# treeClust improves protein co-regulation analysis due to robust selectivity for close linear relationships

**DOI:** 10.1101/578971

**Authors:** Georg Kustatscher, Piotr Grabowski, Juri Rappsilber

**Author notes:** Communicating author.

## Abstract

Gene co-expression analysis is a widespread method to identify the potential biological function of uncharacterised genes. Recent evidence suggests that proteome profiling may provide more accurate results than transcriptome profiling. However, it is unclear which statistical measure is best suited to detect proteins that are co-regulated. We have previously shown that expression similarities calculated using treeClust, an unsupervised machine-learning algorithm, outperformed correlation-based analysis of a large proteomics dataset. The reason for this improvement is unknown. Here we systematically explore the characteristics of treeClust similarities. Leveraging synthetic data, we find that tree-based similarities are exceptionally robust against outliers and detect only close-fitting, linear protein – protein associations. We then use proteomics data to demonstrate that both of these features contribute to the improved performance of treeClust relative to Pearson, Spearman and robust correlation. Our results suggest that, for large proteomics datasets, unsupervised machine-learning algorithms such as treeClust may significantly improve the detection of biologically relevant protein – protein associations relative to correlation metrics.

## INTRODUCTION

Genes with related biological functions tend to be active in the same biological conditions. This is the basis of gene coexpression analysis, a method that predicts the function of unknown genes by comparing their expression profiles to those of well-studied genes (1–5). A typical coexpression study detects gene activity by measuring mRNA abundances of many genes in a range of biological samples or conditions. In a second step, the similarity of expression profiles, i.e. the extent of coexpression between any two genes, is determined by correlation analysis. Finally, pairwise coexpression coefficients are aggregated into a gene coexpression network, through which any uncharacterised genes in the dataset may become associated with clusters of genes of well-defined biological functions.

Many variations of this basic approach have been developed over the past two decades (6). For example, several coexpression measures have been explored as alternatives to Pearson’s correlation, including Spearman’s correlation, Biweight midcorrelation, Mutual Information and simple regression models (7, 8). No single measure appears to be superior for every dataset, as the optimal choice of measure depends on various characteristics of a given dataset, such as the frequency of outliers and missing values.

A recent, fundamental change to the expression profiling setup was made possible by improvements in the field of quantitative proteomics: the use of protein abundances rather than mRNAs as readout for gene activity. This increases the accuracy of gene function prediction, because protein abundances are better indicators of gene function than mRNA levels, at least in human (9–11) and mouse (12). We have recently reported ProteomeHD, a dataset that quantifies the response of 10,323 human proteins to 294 biological perturbations using isotope-labelling mass spectrometry (13) (https://www.biorxiv.org/content/10.1101/582247v1, in revision). ProteomeHD is a heterogeneous dataset, incorporating a wide range of perturbation experiments from different laboratories, such as inhibitor treatments, differentiation time courses and cancer cell line comparisons. We compared different coexpression measures for their ability to detect proteins that are co-regulated in response to these perturbations. Surprisingly, we found that the unsupervised machine-learning algorithm treeClust (14, 15) provided a striking improvement over established correlation-based metrics. However, the reason for this improvement remained unclear, because treeClust is a novel algorithm that works in a fundamentally different way to previously used coexpression metrics.

The treeClust algorithm uses recursive partitioning (16–18) to create decision trees. Such trees are normally used for supervised classification or regression tasks. In contrast, treeClust uses decision trees to calculate a dissimilarity measure in an unsupervised manner. To do so, treeClust first creates a number of decision trees aimed to dissect the dataset by growing one decision tree for each variable in turn, using it as response variable and all remaining variables as predictors. This part of the algorithm is reminiscent of an established approach to impute missing data which uses Random Forests (19). However, in a second step, treeClust calculates dissimilarities between any two observations based on the proportion of trees in which they land in different leaves, using Gower’s distance (20). While treeClust dissimilarities appear to perform well in practical applications (21, 22), some of their basic properties remain unclear, especially in the context of gene expression analysis. For example, do treeClust dissimilarities capture linear or non-linear associations? How is treeClust performance affected by missing data, outlier data and noise? Here, we set out to systematically address these questions and benchmark treeClust performance on both synthetic and real proteomics data.

## EXPERIMENTAL PROCEDURES

### General data analysis and availability

All data processing and analysis has been performed using R version 3.5.1 (23). All data and R scripts required to reproduce the results of this manuscript are available in the following GitHub repository: https://github.com/Rappsilber-Laboratory/treeClust-benchmarking.

The R package data.table (24) was used for fast data processing. Figures were prepared using ggplot2 (25), gridExtra (26), cowplot (27) and viridis (28).

### Generation of synthetic datasets

Synthetic datasets were generated using a custom function in R. The function populates a table with values that are randomly sampled from a normal distribution, but includes a user-specified number of observations that have a defined linear relationship with each other. The following properties of the thus created datasets can be manipulated: number of variables (i.e. samples or experiments), number of observations (i.e. proteins), percentage of protein pairs that should have a linear relationship, percentage of outlier data, percentage of missing values and the extent of scatter around the regression line (i.e. biological or measurement noise). Outlier data points are created by random sampling from a broader normal distribution than the rest of the data.

In addition to positive linear relationships (y ∼ x), we tested relationships that were exponential (y ∼ e^x^), logistic (y ∼ 4 / (1 + e^−5x^)) and quadratic (y ∼ x^2^), as well as linearly anti-correlated (y ∼ −x).

### Real biological datasets

ProteomeHD has been documented in detail elsewhere (13). In short, it is a data matrix consisting of 10,323 proteins whose abundance changes in response to 294 biological perturbations have been determined by quantitative mass spectrometry, using stable isotope labelling by amino acids in cell culture (29). To distinguish between genuine, biologically relevant protein – protein associations (true positives) and likely false positive interactions we used a gold standard based on the Reactome database (30), which was also described previously (13).

### Comparison of coexpression measures

Pearson’s correlation coefficients and Spearman’s rank correlation coefficients were calculated using R base functions (23). Biweight Midcorrelation (bicor) was calculated with default settings using the R package WGCNA (31, 32). TreeClust dissimilarities were calculated using the R package treeClust (14, 15), with the d.num parameter set to 2. When applying treeClust to ProteomeHD rather than synthetic data, we set the rpart complexity parameter to 0.105 and the treeClust serule parameter to 1.8. These settings had been optimised previously for ProteomeHD (13), providing approximately a 10% performance improvement over default values when assessed against the Reactome gold standard.

Performance of coexpression measures was compared by precision – recall (PR) analysis using the R package PRROC (33). True positive (linear or nonlinear) and false positive (random) associations for the PR analyses were known a priori for synthetic data and annotated using the Reactome gold standard for ProteomeHD. To test the impact of various data characteristics, synthetic dataset were generated in triplicate and the result is shown as the average area under the PR curves, with error bars indicating the standard error of the mean. No replicates were used for the combinatorial testing of two dataset characteristics (Fig. 2C, G and H).

### Model fitting in real proteomics data

Base R functions were used to fit and analyse linear models for pairs of proteins in ProteomeHD. Fold-changes of each protein pair were rescaled to fall between 0 and 1 before fitting the model. Outliers were defined as data points with absolute studentized residuals or a Mahalanobis distance larger than 2. Non-linear models were fit using nonlinear least squares. Exponential models (y ∼ a + exp(b)^x^) and logistic models (y ∼ a / (1 + e^−b(x-c)^)) were said to outperform the corresponding linear model (y ∼ a + bx) if their residual sum of squares (RSS) was at least 10% smaller.

## RESULTS AND DISCUSSION

### A coexpression take on Anscombe’s quartet

In the protein co-regulation analysis of ProteomeHD treeClust outperformed common coexpression measures: Pearson’s correlation coefficient (PCC), Spearman’s rank correlation (rho) and Biweight midcorrelation (bicor). To explore possible reasons for this we used Anscombe’s quartet (34) as a starting point. These four 11-point datasets illustrate several key issues that can negatively affect the performance of Pearson’s correlation (Fig. 1A). For example, PCC can falsely identify a linear correlation when there is a non-linear relationship between two variables. In addition, outlier data located far off the regression line can lead to an underestimation of the correlation. Similarly, outliers can cause a high PCC when in fact no correlation between two variables exists at all. Spearman’s rho and bicor are also affected by these issues, albeit to a much lesser extent than PCC (Fig. 1A).

**Figure 1.**
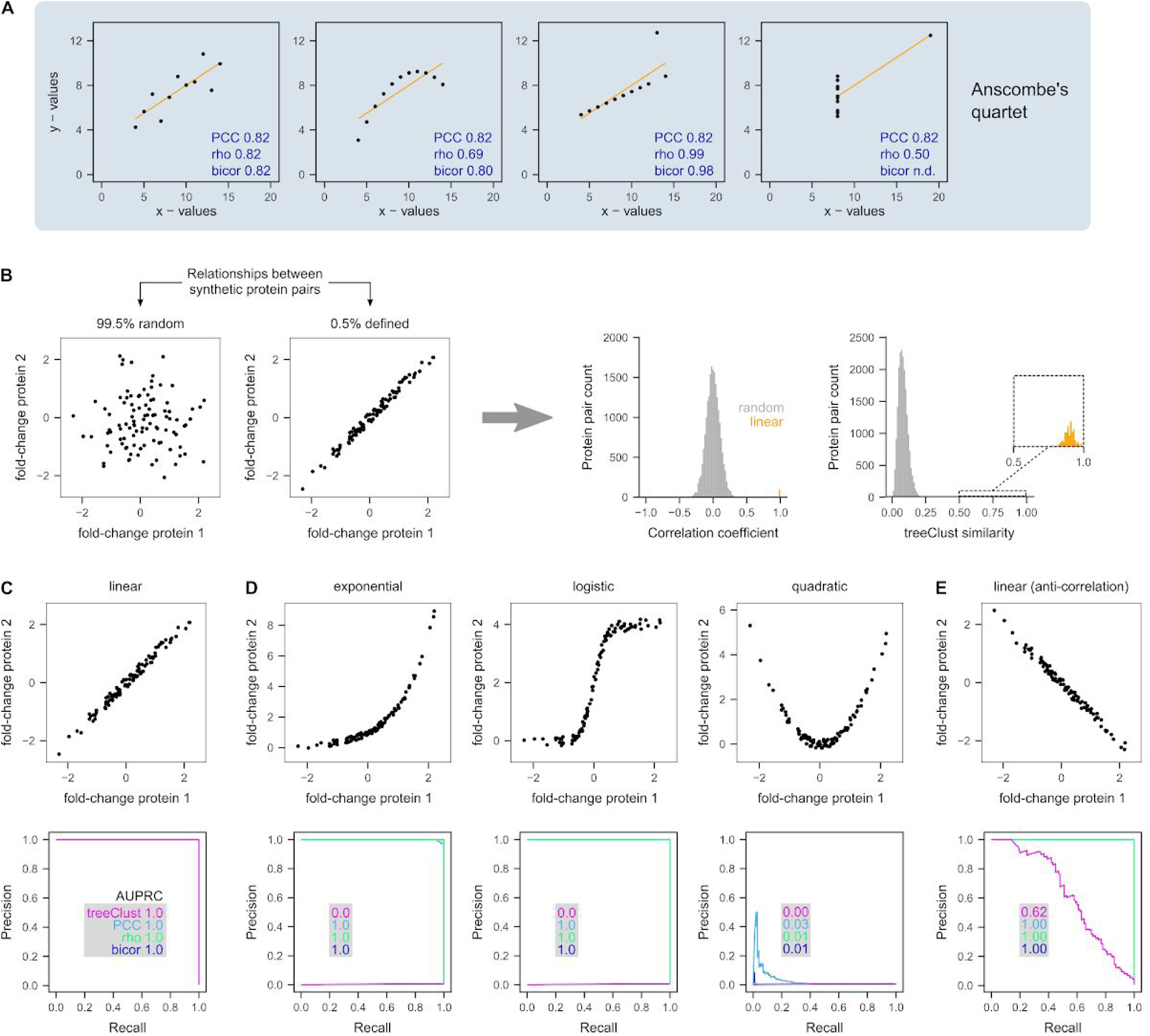
treeClust detects specifically positive linear associations. (**A**) Anscombe’s quartet is a collection of four 11-point datasets that have the same Pearson’s correlation but very different relationships. If these variables were genes and the values were expression measurements, PCC would significantly underestimate (example 2 and 3) or overestimate (example 4) the extent of their coexpression. (**B**) To test how treeClust deals with such data we created a synthetic dataset consisting of 100 variables and 200 proteins. The dataset is designed such that out of all possible 19,900 combinations between these proteins, 0.5% have a defined relationship while the remaining 99.5% of pairs have not. Histograms show the resulting distribution of correlation coefficients or treeClust similarities. In this best-case scenario a complete separation between random and defined pairs is easily achieved. (**C**) Precision – recall (PR) analyses show that treeClust separates linear from random relationships perfectly, resulting in an area under the PR curve (AUPRC) of 1. The same result is observed for the three tested correlation-based metrics: PCC, Spearman’s rho and biweight midcorrelation (bicor). The four PR curves overlap fully. (**D**) TreeClust completely fails to detect exponential or logistic relationships (AUPRC = 0). In contrast, these pairs still score high enough with PCC, rho and bicor to be completely separated from the pool of random associations. No metric detects quadratic relationships. (**E**) Anti-correlations are not identified well by treeClust.

**Figure 2.**
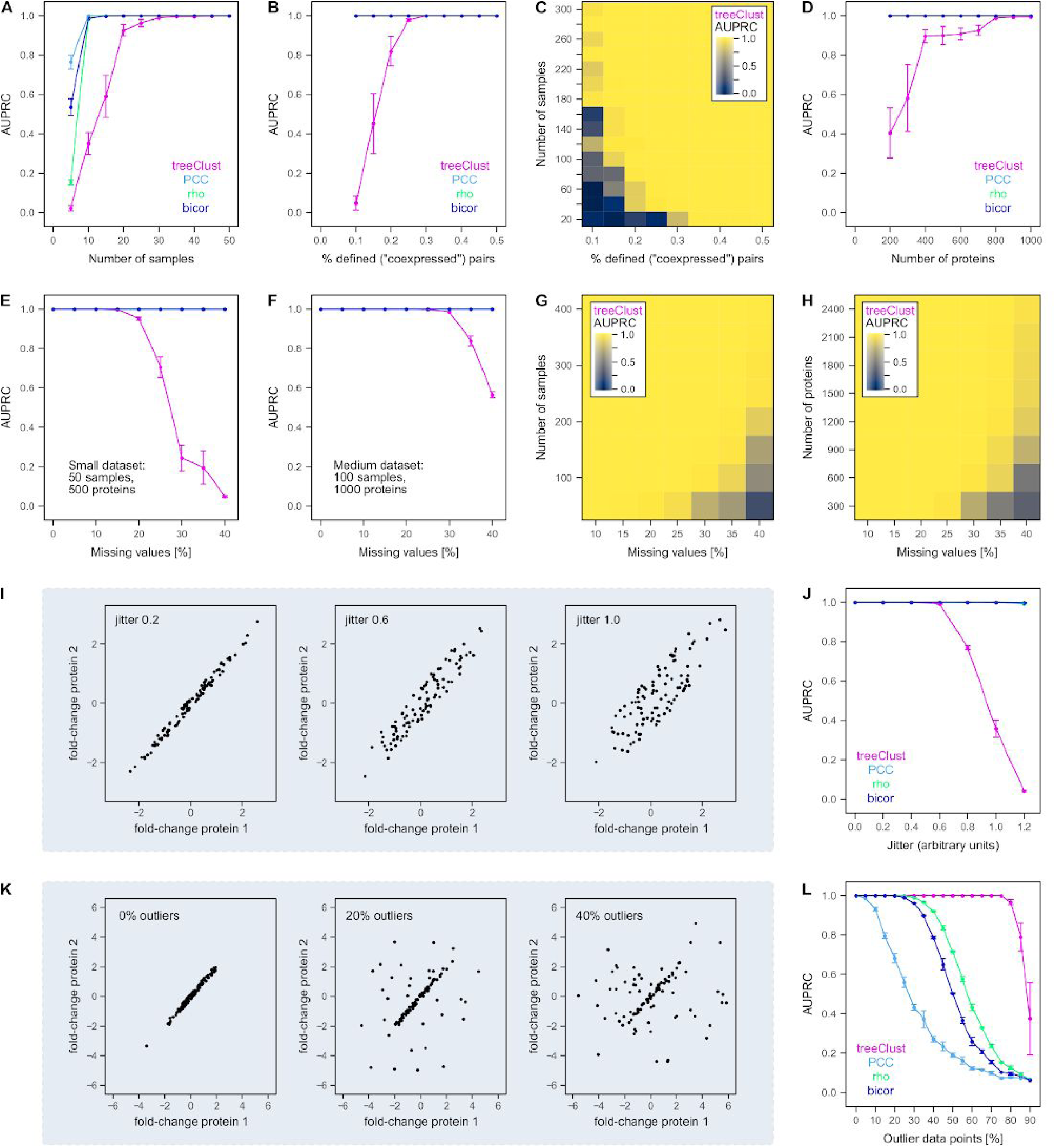
Using synthetic data to benchmark treeClust performance. (**A**) Increasing the number of samples (variables) in a synthetic dataset improves the performance of treeClust, Pearson’s correlation (PCC). Spearman’s correlation (rho) and Biweight Midcorrelation (bicor). TreeClust needs more samples for optimal performance than the correlation metrics. Each synthetic dataset, containing 500 proteins and 0.3% linear relationships, was created in triplicate. Points show the average area under the precision recall curve (AUPRC) obtained for each setting. Error bars show the standard error of the mean. (**B**) Same as A but increasing the linear associations in datasets of 50 samples and 500 proteins. This has no impact on the three correlation metrics, so their curves overlap fully at AUPRC 1. (**C**) Combinatorial impact of the two parameters on treeClust AUPRC is shown through a colour-gradient. N proteins = 500. (D) Same as A but increasing the number of observations (proteins). N samples = 20, 0.3% linear associations. (**E**, **F**) Adding missing values to a small and medium dataset, respectively. (**G**) Combinatorial impact of missing values and sample number on treeClust performance. N proteins = 1,000. (**H**) Same as G but modifying the number of proteins (N samples = 150). (**I**) Scatterplots illustrating the effect of increasing the difference between variables. (**J**) This increasing dispersion strongly impairs detection of linear associations by treeClust, but not correlation metrics. (**K**) Scatterplots illustrating the effect of adding outlier data points. (**L**) Impact of outlier data on the four coexpression measures. TreeClust is exceptionally robust against outliers.

We then asked how treeClust deals with Anscombe’s quartet. However, it is not possible to simply calculate treeClust dissimilarities for Anscombe’s four variable pairs. This is because, being a machine-learning algorithm, treeClust requires an input dataset with many variables in order to build informative decision trees. Therefore, we created a series of synthetic datasets that allow us to systematically assess the properties of treeClust dissimilarities and compare them to the properties of common coexpression measures. For example, we created a synthetic dataset consisting of 100 variables (experiments, samples or biological conditions) and 200 observations (proteins). The dataset is built in such a way that 99.5% of the resulting pairwise “protein – protein” associations are random, i.e. values for both proteins are random samples of a normal distribution (Fig. 1B). The remaining 0.5% pairs are designed to have a clearly defined, linear relationship across the 100 “experiments”. These pairs have a PCC close to 1, which clearly sets them apart from the distribution of the random pairs (Fig. 1B). On such a synthetic dataset, treeClust generates 100 decision trees that assign very different dissimilarities to random and linear associations (Fig. 1B). Indeed, in this simple best-case scenario, all of the four tested coexpression measures separate random and defined pairs perfectly, resulting in precision – recall curves with an area of 1 (Fig. 1C).

Note that in this manuscript we show treeClust similarities rather than dissimilarities (similarity = 1 – dissimilarity), in order to make the comparison with correlation metrics more intuitive (Fig. 1B).

### Linear vs non-linear relationships

We then proceeded to modify various properties of the synthetic datasets and assessed how they affect treeClust. First, we asked which types of associations are detected by treeClust. For example, we replaced the 0.5% linearly correlated pairs with exponential relationships (Fig. 1D). This does not affect Pearson, Spearman or robust correlation, which still yield an area under the precision – recall curve (AUPRC) of 1. Although exponentially related pairs receive lower correlation coefficients than linear ones, their coefficients are still much higher than those of random pairs. Surprisingly and in stark contrast to the correlation measures, treeClust does not detect exponential relationships at all, yielding an AUPRC of 0 (Fig. 1D). We obtained the same result for logistic relationships (Fig. 1D). None of the coexpression measures detects quadratic relationships. Finally, we tested if treeClust detects negative linear associations, i.e. anti-correlation. We find that treeClust only partially separates anti-correlated from random associations, suggesting that low treeClust similarities indicate a lack of correlation rather than anti-correlation (Fig. 1E).

We conclude that, in the conditions tested here, treeClust specifically captures positive linear “protein – protein” associations. This property could be explained by the fact that treeClust dissimilarities reflect how often two observations land in the same decision tree leaf. A split in a decision tree is less likely to separate two linearly associated proteins than exponentially related or anti-correlated proteins. For example, if protein X1 is upregulated 1.5-fold in a given experiment, a linearly related protein X2 may be upregulated 1.6-fold. These proteins would only land in different leaves if a split occurred between 1.5 and 1.6. However, if protein X2 was exponentially related to X1 it may be upregulated by 4.6-fold, increasing the margin within which a split could occur such that the two proteins land in different leaves. Similarly, if two proteins are anticorrelated they rarely end up in the same leaf and thus cannot be flagged up by treeClust as being linked.

### Size and structure of the dataset

We next investigated how basic data characteristics such as the number of variables and observations affect treeClust. In principle, treeClust performance is expected to improve with the amount of data it is presented with, because more data may allow treeClust to build more informative decision trees and thus learn better to distinguish between random and genuine linear associations. To test this, we constructed a series of synthetic datasets with increasing dimensions.

First, we increase the number of variables, i.e. samples (Fig. 2A). Under our test conditions treeClust requires around 40 samples to reach optimal performance, i.e. an AUPRC of 1. In contrast, the three correlation metrics only need ∼15 samples to reliably identify all genuinely correlated proteins (Fig. 2A). A likely explanation for this difference is that treeClust builds one decision tree per sample, so increasing the number of variables also increases the number of decision trees and thus the reliability of the resulting dissimilarities.

Next, we modify the percentage of linear associations (i.e. coexpressed proteins) in the dataset (Fig. 2B). This has no impact on the correlation measures but affects treeClust performance, suggesting that the more genuine associations are present in the data the easier it is for treeClust to learn to identify them. Notably, these two data size characteristics are interdependent: increasing the number of samples compensates for a smaller percentage of defined associations and vice versa (Fig. 2C).

Third, we test the impact of having more observations (proteins) in a dataset. We find that increasing the number of proteins to > 1,000 is sufficient for optimal treeClust performance even if only 20 samples are available (Fig. 2D). Larger and more complex input data generally improve the performance of machine-learning algorithms. While increasing observations will not affect the number of decision trees, the increased complexity of the input data allows treeClust to create more informative trees.

In summary, these results show that optimal treeClust performance requires a dataset of a certain size and structure, for example ≥ 50 variables, ≥ 1000 proteins and ≥ 0.4% linear associations. For smaller datasets, for example in the range of 20 variables and 500 observations, traditional correlation-based measures may be better suited for coexpression analysis.

### Missing values

Proteomics data often contain a large number of missing values. In this case, correlation metrics simply focus on those variables that have been measured for both proteins. The decision trees used by treeClust handle missing values through surrogate splits (14, 17), an approach that is generally considered to be sensible only if missing values are sparse. To evaluate the impact of missing values on treeClust performance we randomly introduce missing values in synthetic data. In a dataset with 50 variables and 500 observations, introducing 10% – 15% of missing values has no ill-effect on treeClust performance (Fig. 2E). Beyond that, missing values quickly become detrimental for treeClust. In contrast, they do not pose a problem for correlation metrics as long as a sufficient number of common pairwise measurements remain available (Fig. 2E). However, we find that the impact of missing values depends on the overall dimensions of the input data. For example, with a dataset of 100 samples and 1000 proteins treeClust can already tolerate 20% missing values (Fig. 2F). Consequently, we systematically explore the impact of missing values depending on the number of samples or proteins. We observe that for large datasets treeClust performance does not decrease even if 40% of all values are missing (Fig. 2G, H).

### Goodness-of-fit

We then asked how “tight” coexpression of two hypothetical proteins needs to be for treeClust to detect it. To this end, we increased the dispersion / scatter of values around the linear associations (Fig. 2I). Within the range of parameters tested, increasing dispersion had no appreciable effect on the performance of the three correlation metrics (Fig. 2J). In contrast, treeClust detects only very close, well-fitting linear associations. As for non-linear relationships, the explanation for this behaviour may lie in the likelihood of decision tree splits occurring between two observations. A larger scatter around the regression lines signifies a larger difference between two proteins and therefore an increased probability for them to land in different leaves.

### Outlier data points

Finally, we assessed the impact of outlier data on treeClust dissimilarities. Introducing outlier data points in a synthetic dataset confirms the well known error-proneness of Pearson’s correlation in the presence of outliers (Fig. 2K, L). As expected from Anscombe’s examples, the AUPRC for Pearson’s correlation is halved if around 25% of the measurements are outliers. Spearman and Biweight midcorrelation, which are less susceptible to outliers, handle this level of outliers in our test set without performance decrease (Fig. 2K, L). However, treeClust is exceptionally robust against outlier measurements, even in comparison to Spearman’s rho and bicor. In the synthetic dataset, treeClust performance is completely unaffected by up to 75% outliers. Therefore, treeClust can detect an association between two synthetic proteins if only 25% of the actual measurements show a strong linear relationship.

### Applying the lessons from synthetic data to real proteomics experiments

The synthetic datasets revealed several marked differences between treeClust and traditional correlation-based coexpression measures. One potential disadvantage of treeClust dissimilarities is that they can only be calculated accurately for datasets that fulfill certain requirements on size and structure, including the number of experiments, proteins, percentage of coexpressed protein pairs and missing values.

A dataset like ProteomeHD is well within the margins of optimal treeClust performance identified by the synthetic data. We applied treeClust to 5,013 proteins in ProteomeHD that had been observed in at least a third of the 294 samples. This subset of ProteomeHD contains 35% missing values and a sufficient percentage of genuine linear protein-protein associations. The latter is estimated based on the observation that 3% of all protein pairs in this dataset have strong and significant Pearson’s correlations (PCC > 0.5, Bonferroni adjusted *p-values* < 1e-6). This is well in excess of the 0.5% margin determined on synthetic data, even after accounting for a reasonable fraction of potential false-positives.

Using the synthetic data we identified two potential reasons for the improved co-regulation analysis of ProteomeHD. First, treeClust detects exclusively close, linear relationships, and this selectivity may make it better suited to detect genuine biologically relevant associations. Second, treeClust is exceptionally robust towards outlier measurements. Next, we tested which of these may be relevant for ProteomeHD.

### Outliers in ProteomeHD

We first evaluated the impact of outlier measurements in ProteomeHD. We used two different statistical methods to detect outliers, as they identify distinct types of outliers. Data points with large studentized residuals are regression outliers, meaning they are far from the regression line but not necessarily unusual with regards to the overall distribution of the ratios (Fig. 3A). This type of outlier may lead to an underestimation of the real correlation coefficient. In contrast, outliers with a large Mahalanobis distance are far from the bulk of the data but can be close to the regression line (Fig. 3A). These outliers may lead to an overestimation of the real correlation coefficient. In principle, regression and Mahalanobis outliers could be seen as real-life examples of the outliers shown in Anscombe’s third and fourth dataset, respectively.

**Figure 3.**
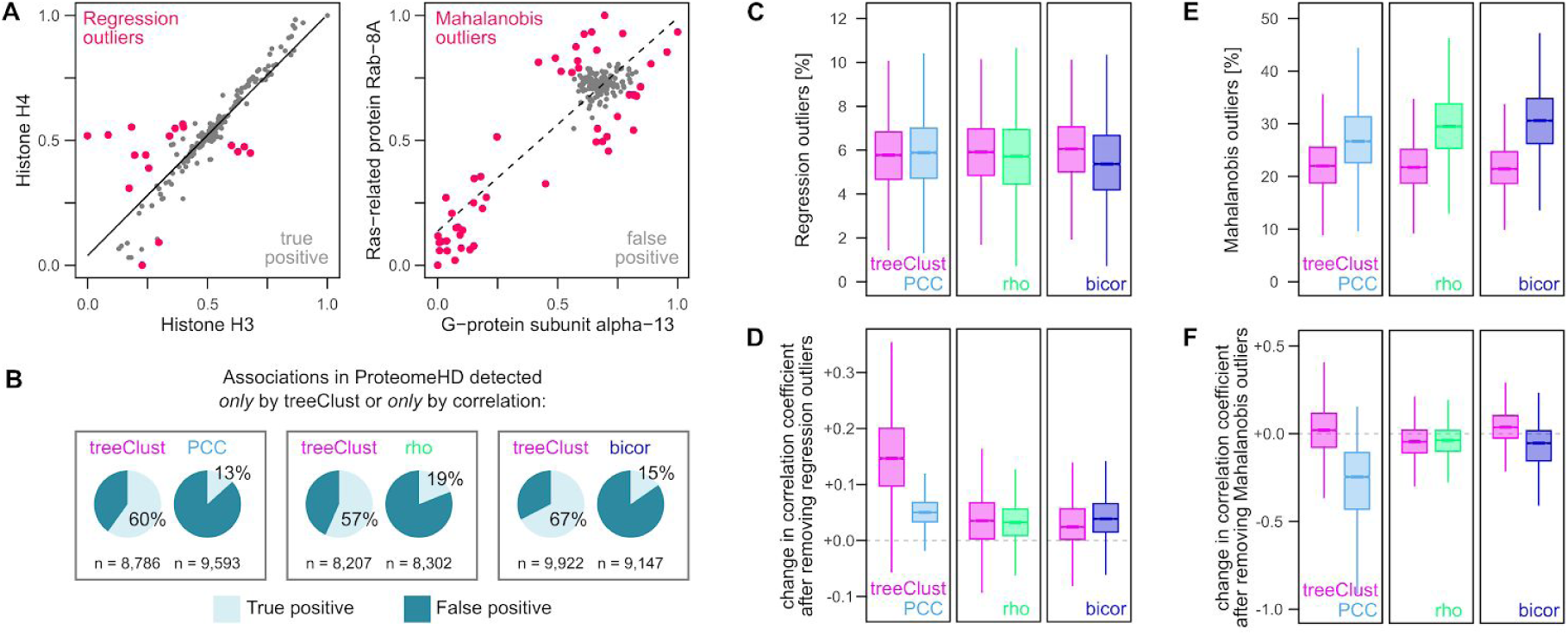
Outliers in ProteomeHD and their impact on correlation metrics. (**A**) Two examples protein pairs from ProteomeHD illustrate the two different types of outliers. Regression outliers are detected via studentized residuals and are located far away from the regression line. These outliers will decrease correlation coefficients. Outliers detected via their Mahalanobis distance are located far away from the bulk of the data, but can be close to the regression line. As in the example show, these outliers can cause high correlation cofficients even if no general correlation exists. Fold-changes have been scaled to lie between 0 and 1. (**B**) Co-regulated protein pairs were divided into those detected by treeClust but not by PCC and vice versa. Separate comparisons were made for pairs detected by treeClust but not rho, and treeClust but not bicor. The pairs in the resulting groups were annotated using Reactome into known, biologically relevant interactions (true positives) and pairs that were unlikely to have any biological associations (false positives). Note that treeClust-specific pairs tend to be true positives, whereas correlation-specific pairs tend to be false positives. (**C**) The number of regression outliers is very similar in all six groups. (**D**) Removing regression outliers increases the PCC of protein pairs that were previously detected only by treeClust, suggesting PCC missed some of these pairs because of regression outliers. This is not the case for pairs missed by rho or bicor. (**E**) Mahalanobis outliers are more frequent in protein pairs detected by all three correlation metrics than pairs specific to treeClust. (**F**) Removing Mahalanobis outliers decreases the PCC of pairs that were originally detected only by PCC, suggesting their PCC was supported mainly by the Mahalanobis outliers. The correlation of pairs detected only by rho or bicor is not affected strongly by removing Mahalanobis outliers.

We then tested if either of these outlier types explains why treeClust outperforms PCC for ProteomeHD data. For this we compared protein pairs that scored high (i.e. ranking in the top 0.1% pairs) with one method but were not detected (i.e. not ranked in top 0.5% pairs) by the other. This resulted in the following two groups of protein pairs: (a) 8,786 protein pairs with high treeClust similarities that were not detected as co-regulated by their PCCs; (b) 9,593 protein pairs with high PCCs that were not detected by treeClust. Functional annotation of these groups using a gold standard based on Reactome revealed that 60% of protein pairs found exclusively by treeClust are known true-positive associations, compared to only 13% of the PCC-specific pairs (Fig. 3B). Therefore, treeClust-specific co-regulation pairs are predominantly true interactions missed by PCC, whereas PCC-specific pairs are mostly unrelated proteins falsely identified as co-regulated by PCC. Similar distributions were observed when comparing treeClust to Spearman’s rho and bicor (Fig. 3B).

Next, we asked if regression outliers in ProteomeHD may explain the difference between these groups. Surprisingly, we find that the number of regression outliers is very similar for treeClust-specific and PCC-specific protein pairs (5.8% vs 5.9% on average), as well as treeClust and rho-specific and bicor-specific pairs (Fig. 3C). However, the impact of outliers may not just stem from their number but also from their actual position and distribution compared to the rest of the data. We therefore removed outliers and measured the effect this had on the correlation coefficients. Indeed, removing regression outliers has a strong impact on co-regulation pairs that had been detected by treeClust but not by PCC, increasing their average PCC by 0.15 (Fig. 3D). This suggests that treeClust-specific co-regulation pairs tend to be genuine, biologically relevant interactions that are missed by PCC due to regression outliers. However, removing regression outliers had no dramatic effect on pairs that had been missed by rho or bicor (Fig. 3D).

In contrast to regression outliers, Mahalanobis outliers are clearly enriched among pairs only detected by PCC (27% vs 22% on average), or only by rho or bicor (Fig. 3E). Removing Mahalanobis outliers has a striking impact on PCC-specific pairs, reducing their average PCC by −0.29 (Fig. 3F). This indicates that co-regulated pairs detected by PCC – but not treeClust – are predominantly false-positive interactions whose high PCC is driven by Mahalanobis-type outliers. Associations detected only by rho or bicor, although enriched for Mahalanobis outliers, do not lose their high correlation coefficients by removing these outliers (Fig. 3F).

In summary, these results suggest that outliers are a key factor explaining why treeClust outperforms PCC in the analysis of ProteomeHD data. However, its improvement over rho and bicor is unlikely to be due to better outlier handling.

### Goodness-of-fit of genuine associations in ProteomeHD

Increasing the scatter of values around the regression line led to a dramatic reduction of treeClust similarity in synthetic data (Fig. 2J). To quantify the overall difference between two proteins in real biological data we use the mean absolute error (MAE). Two protein pairs with very similar correlation coefficients can have vastly different MAEs (Fig. 4A). As expected, of the example pairs shown in Fig. 4A, the pair with the small MAE reflects a genuine biological association and receives a high treeClust similarity. In contrast, the pair with the large MAE is composed of two unrelated proteins and is not detected as co-regulated by treeClust (Fig. 4A). This suggests that treeClust may distinguish better than correlation measures between close, real interactions and loose, biologically irrelevant trends.

**Figure 4.**
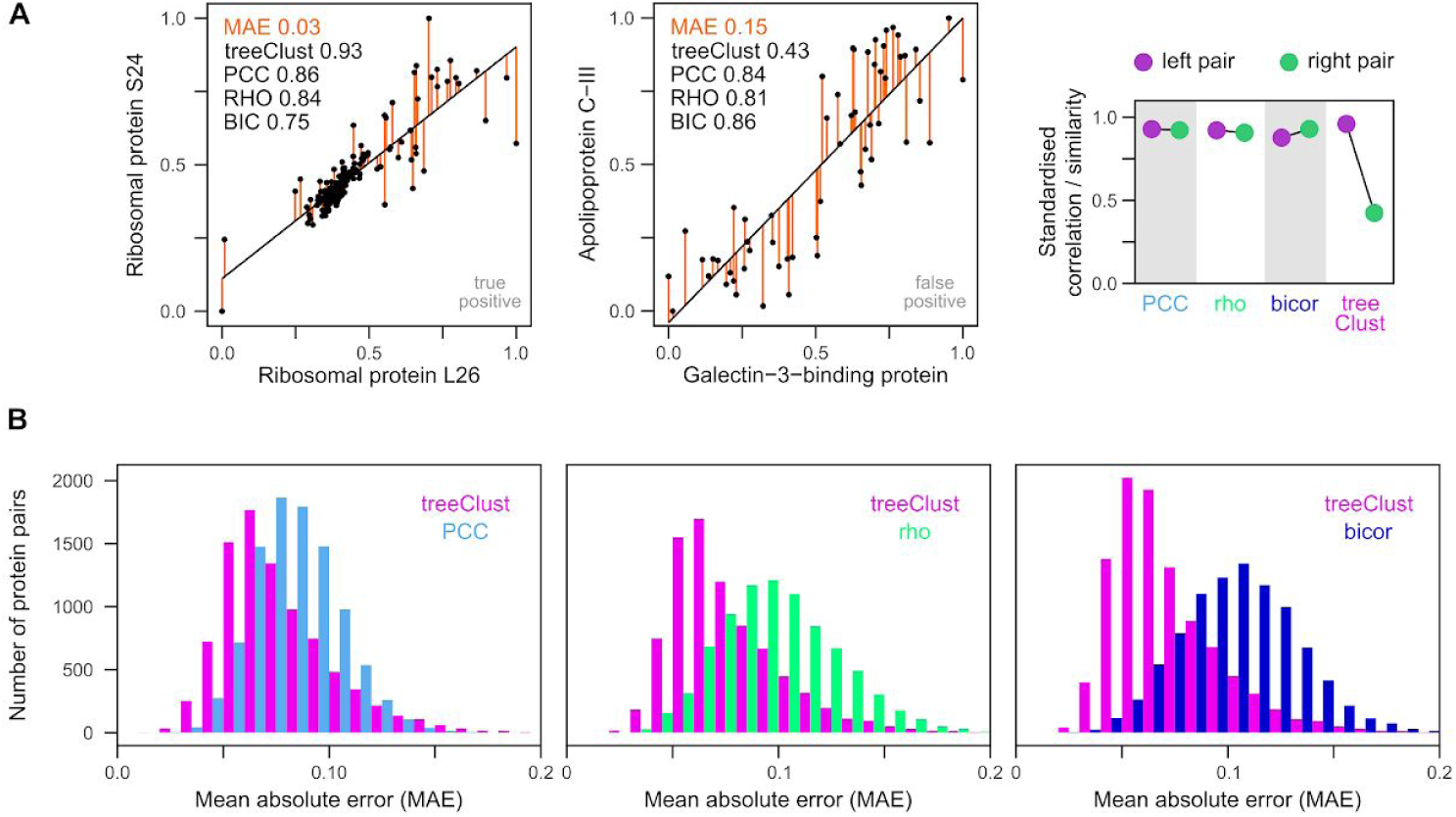
Goodness-of-fit is a critical parameter for detecting genuine associations in ProteomeHD. (**A**) Two examples pairs from ProteomeHD to illustrate close and lose – fitting regression. Both pairs have similar regression slopes and correlation coefficients. However, the left pair has a much smaller mean absolute error (MAE). MAE is the average of the absolute residuals (orange). The left pair is a known, biologically relevant interaction documented by the Reactome gold standard, the right pair is not. Unlike the correlation metrics, treeClust assigns the right pair are much weaker similarity. The difference between the left and right example pair is shown by the scatterplot on the right. For this plot the correlation metrics and treeClust similarities were standardised to fall within a range of [0,1] to make them comparable. (**B**) Systematic comparison of MAEs from protein pairs that scored high both with treeClust and PCC (or rho, or bicor), or pairs that scored high with either metric alone. Protein pairs exclusively detected by correlation metrics tend to have much higher MAEs, possibly explaining why they are predominantly false-positive hits.

To assess this possibility in a systematic way we analysed the MAE distribution of all protein pairs that receive high correlation coefficients but low treeClust similarities, and vice versa. We find that protein pairs exclusively detected by rho or bicor tend to have much higher MAEs than those exclusively detected by treeClust (Fig. 4B). Interestingly, the difference in MAE distribution is not as pronounced between treeClust and PCC. Taken together this suggests that treeClust outperforms PCC mainly due to its outlier handling, whereas its improvement over rho and bicor is predominantly due to treeClust taking into account the “goodness-of-fit” of an association.

### Lack of non-linear relationships in ProteomeHD

The selectivity of treeClust for linear relationships implies that it may fail to detect non-linear relationships that may be biologically relevant. We therefore investigated whether any genuine non-linear protein – protein associations exist in ProteomeHD. For this we fitted linear, exponential and logistic models to the correlation- or treeClust-specific protein pairs. For each pair we then select the best-fitting model based on the residual sum of squares (RSS). We find that exponential models rarely fit better than the linear regression models, but surprisingly, logistic models often do (Supplementary Fig. S1A). However, closer inspection of the data reveals that these cases are not genuine exponential or sigmoid relationships (Supplementary Fig. S1B). In contrast, the improved fit of the non-linear models is driven by Mahalanobis-type outliers. Removing these outliers also drastically decreases the number of instances in which non-linear models fit better than linear ones (Supplementary Fig. S1A). In summary, we have not been able to identify any clear non-linear relationships in ProteomeHD.

## CONCLUSION

Having found treeClust to be a powerful alternative to correlation metrics for the detection of protein – protein links in proteomics data recently (13) we here demonstrated possible reasons for this observation. treeClust is exceptionally robust against outliers and only identifies close-fitting, positive linear associations. In real proteomics datasets, these type of associations appear to be the biologically most relevant ones. Obvious disadvantages of using unsupervised machine-learning for this task are the required size and composition of the input data. At the moment, few proteomics datasets exists that are large enough, i.e. covering hundreds of conditions for thousands of genes, while maintaining a sufficiently low percentage of missing values for treeClust to work effectively. This applies to ProteomeHD and is likely going to become more prevalent in the near future, also thanks to the efforts of the ProteomeXchange consortium (35, 36). However, also smaller datasets can be analysed, by applying the algorithm many times to different subsets of the data and collecting the average similarities across these models (a bootstrapping approach) (13). It will be interesting to see how treeClust fares with other omics data types and other application areas where currently correlation approaches are in use.

## ACKNOWLEDGEMENTS

This work was supported by the Wellcome Trust through a Senior Research Fellowship to J.R. (grant number 103139). The Wellcome Centre for Cell Biology is supported by core funding from the Wellcome Trust (grant number 203149).

---

**Supplementary Figure S1.**
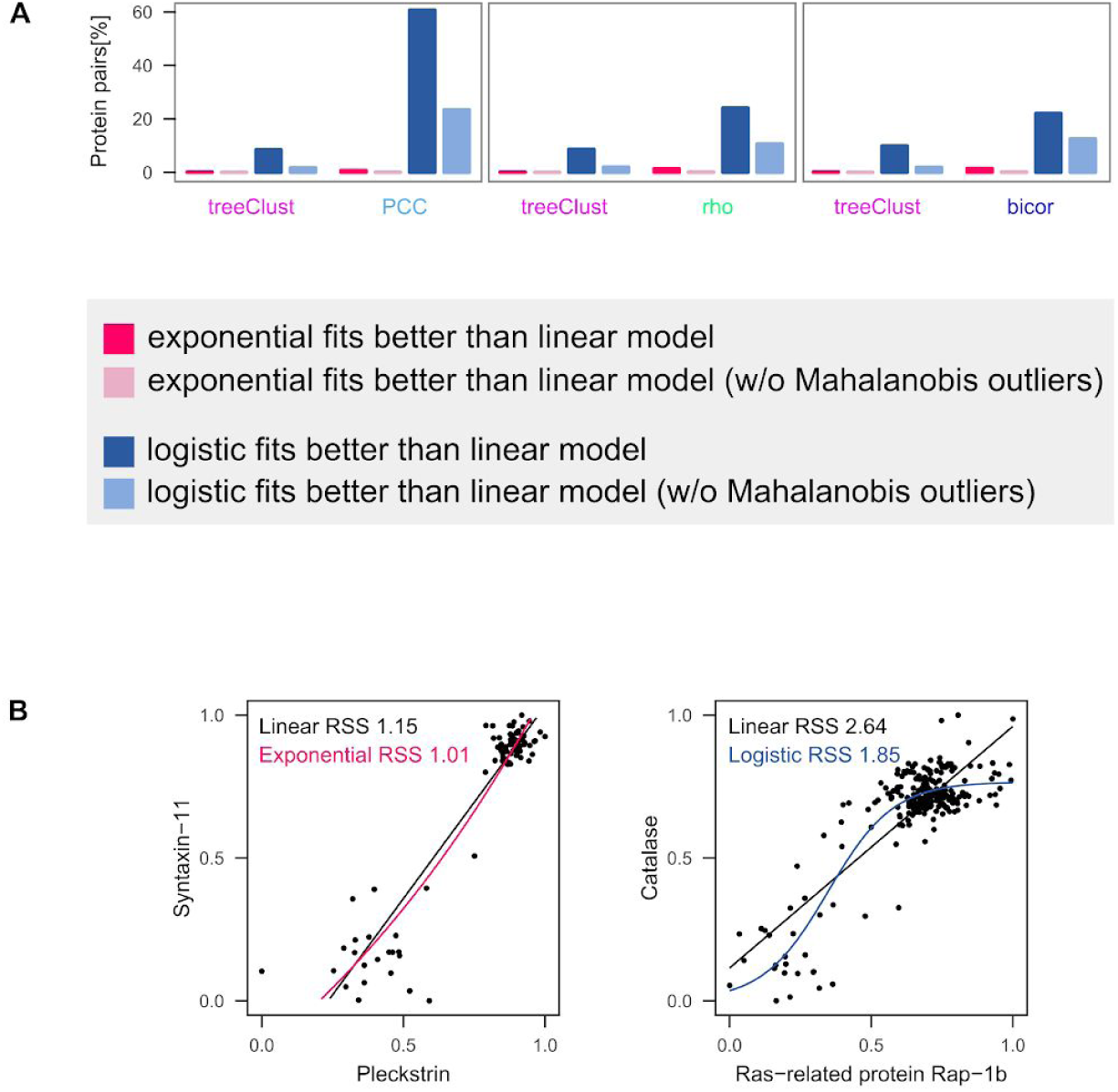
Lack of genuine non-linear relationships in ProteomeHD. (**A**) Exponential and logistic (sigmoid) models were fitted to all protein pairs that scored high with treeClust or the three correlation metrics. Model fit was compared through their residual sum of squares (RSS). Exponential models only fitted better than linear ones in rare cases, but logistic models often did. Around half of the protein pairs detected specifically by PCC are better explained by a logistic than a linear model. However, this is mainly driven by Mahalanobis-type outliers. Removing those strongly reduces the number of logistic models outfitting the linear ones. (**B**) Two example regressions where an exponential (left) or logistic (right) model fits better than a linear one. Note that this clearly reflects overfitting due to outliers rather than genuine non-linear relationships.

